# Gallic Acid Disrupts Aβ_1-42_ Aggregation and Rescues Cognitive Decline of APP/PS1 Transgenic Mouse

**DOI:** 10.1101/258848

**Authors:** Mei Yu, Xuwei Chen, Jihong Liu, Quan Ma, Hao Chen, Lin Zhou, Sen Yang, Lifeng Zheng, Chengqing Ning, Jing Xu, Tianming Gao, Sheng Tao Hou

## Abstract

Alzheimer’s disease (AD) treatment represents one of the largest unmet medical needs. Developing drugs capable of preventing Aβ aggregation is an excellent approach to prevent and treat AD. Here, we show that gallic acid (GA), a naturally occurring polyphenolic small molecule rich in grape seeds and fruits, has the capacity to alleviate cognitive decline of APP/PS1 transgenic mouse through reduction of Aβ_1-42_ aggregation and neurotoxicity. Oral administration of GA not only improved the spatial reference memory and spatial working memory of early stage AD mice (4-month-old), but also significantly reduced the more severe deficits in spatial learning, reference memory, short-term recognition and spatial working memory of the late stage AD mice (9-month-old). The hippocampal long-term-potentiation (LTP) was also significantly elevated in the GA-treated late stage APP/PS1 AD mice. Atomic force microscopy (AFM), dynamic light scattering (DLS) and thioflavin T (ThT) fluorescence densitometry analyses showed that GA can reduce Aβ_1-42_ aggregation from forming toxic oligomers and fibrils. Indeed, pre-incubating GA with oligomeric Aβ_1-42_ reduced Aβ _1-42_-mediated intracellular calcium influx and neurotoxicity. Molecular docking studies identified that the 3,4,5-hydroxyle groups of GA were essential in noncovalently stabilizing GA binding to the Lys28-Ala42 salt bridge and the -COOH group is critical for disrupting the salt bridge of Aβ_1-42_. The predicated covalent interaction through Schiff-base formation between the carbonyl group of the oxidized product and ε-amino group of Lys16 is also critical for the disruption of Aβ_1-42_ S-shaped triple-β-motif and toxicity. Together, these studies demonstrated that GA can prevent and protect the AD brain through disrupting Aβ_1-42_ aggregation.

## Introduction

Alzheimer’s disease (AD) is the most common form of senile dementia affecting millions of people worldwide. Characterized by the gradual loss of cognitive functions due to abnormal deposition of extracellular fibrillar amyloid beta peptides (Aβ) and intracellular neurofibrillary tangles in the brain, AD poses tremendous social and economic burdens for families and societies ^1,2^. It has been hypothesized that targeting Aβ neurotoxicity is a potentially effective method to treat AD. Unfortunately, after decades of intensive pursuit, there is still no immediate hope of translating such knowledge into a pharmaceutical therapy for AD ^3,4^. For example, cholinesterase inhibitors were the first generation of such candidate therapeutics ^5–7^. A second generation of candidate therapeutics includes attempts to inhibit the formation of amyloid plaques. Active and passive immunization approaches are presently being developed ^8^, despite of the adverse immune reactions in the active human immunization trials in some patients ^9,10^. Another alternative has been to reduce the levels of Aβ in the brain by inhibiting the proteolytic enzymes associated with trimming APP into Aβ 40 or 42 ^9^.

The Aβ_1-42_ fibril is the predominant constituent of amyloid plaques despite Aβ_1-40_ being more abundant in the plasma ^11^. The observation that Aβ_1-42_ oligomers form calcium conducting channels in the neuronal bilayer membranes ^12,13^ led to the search for calcium channel inhibitors. The nonspecific Aβ channel blockers, such as tromethamine (Tris) and Zn^2+^, have been reported to inhibit Aβ neurotoxicity ^14,15^. Glutamate receptors, including NMDA type and AMPA type receptors, are also involved in Aβ neurotoxicity through increased Ca^2+^ influx ^13^. Thus, calcium homeostasis plays critical roles in the mechanisms of Aβ-induced neurotoxicity ^16,17^. Drugs like memantine targeting glutamate receptor-mediated intracellular Ca^2^+ influx have been developed to provide neuroprotection to treat mid- to late-stage AD ^18,19^.

An entirely different approach to Aβ neurotoxicity can be developed is to prevent the formation of oligomerization of Aβ_1-42_. Oligomeric Aβ_1-42_ are the most toxic to the functionality of neurons ^20^. One such molecule is gallic acid (GA, 3, 4, 5-trihydroxybenzoic acid), a naturally occurring polyphenol small molecule exists in many fruits. Past research demonstrated that GA can inhibit fibril formation of the amyloid-beta peptide and kappa-casein, a milk protein which forms amyloid fibrils spontaneously under physiological conditions ^21^. GA also has a variety of other pharmacological activities, including anti-oxidation, scavenging free radicals, anti-inflammation, antimicrobial and anti-cancer activities ^22–24^. Importantly, the anti-oxidation and anti-inflammation properties of GA contribute to neuroprotection against Alzheimer’s disease ^22,25^. Together these studies led us to hypothesize that GA is a potential candidate for AD drug development.

In the present study, APPswe/PS1dE9 double transgenic mouse AD model (APP/PS1) was used to investigate the therapeutic effects of GA in both early and late stages of disease development. The APP/PS1 mouse was generated by co-injecting two vectors encoding the human mutant APP and PSEN1 to produce elevated Aβ levels ^26^. At the age of 4-month-old, these mice begin to develop small amount of Aβ plaque deposition and show working memory deficits ^27^. At the age of 9-month-old, these mice develop pronounced cognitive deficits. A battery of behavioral tests was performed to show that GA was effective in rescuing cognitive deficits during both the early and late stages of AD development. The possible mechanisms were through disrupting Aβ_1-42_ fibril formation. Collectively, these studies demonstrated that GA is a good lead for the development as a therapeutic drug against AD.

## Materials and Methods

### Preparation of Aβ_1-42_

Synthetic human Aβ_1–42_ (>95%) was purchased from GL Biochem (Shanghai, China) and prepared as preciously described ^28^. Briefly, fresh Aβ_1–42_ powder was dissolved in 1,1,1,3,3,3-Hexafluoro-2-propanol (HFIP, Fluka, USA) to achieve a final concentration of 1 mM. The solution was left in quiescence for 1 h at room temperature, which was then distributed into aliquots in new centrifuge tubes with opened lids for evaporation at room temperature for about 3 h. Afterwards, it was followed by drying in a speed-vac concentrator (CentriVap^®^ Concentrator, Labconco, USA) at 37°C for 1h. The lyophilized peptides were stored at -20°C until use. To obtain oligomeric Aβ_1–42_, the HFIP-treated peptides was re-dissolved in DMSO to a concentration of 2.5 mM and sonicated in a water bath for 5 min. The solution was then diluted in Dulbecco’s Modified Eagle Medium/Nutrient Mixture F-12 to 100 μM and incubated for 48 h at 4 °C. After that, the solution was centrifuged at 12000 rpm for 10 min at 4 °C, and the supernatant was collected for the subsequent experiment.

To produce aggregated Aβ_1-42_ ^20^, the HFIP-treated Aβ_1-42_ peptides was re-dissolved in 10 mM NaOH to a concentration of 2.5 mM and sonicated in a cold water bath for 30 min, followed by dilution in PBS to the required concentration. The solution was incubated for 3 days at 37 °C to form aggregated Aβ_1-4_2.

### Neuronal viability assay using CCK8 kit

Neuronal viability was assessed with a cell viability counting kit – 8 purchased from BBI Life Sciences (Shanghai, China). Briefly, primary cortical neurons treated with Aβ_1-42_ for the specified period of time and the amount were collected following the manufacturer’s protocol. Exactly 100 μL of cell suspension was transferred to a 96-well plate at a final concentration of 500,000 cells/mL. Cells were incubated with 10 μL CCK-8 solution from the kit per well for 4 h and assessed with a Perkin-Elmer spectrometer (EnSpire, Perkin-Elmer, USA) at the absorbance of 450 nm.

### Quantification of Aβ plaques using Thioflavin S staining

Thioflavin S staining of frozen brain sections was performed as previously described with minor modifications ^28^. Briefly, mouse brain was fixed in 4% paraformaldehyde for 48 h and dehydrated in 10% and 30% sucrose solution. Coronal sectioning at 10 μm thickness of mouse brain with 10 serial sections was performed in a cryostat. Brain sections were dried in the air and dehydrated in ethanol solution. Brain sections were incubated with 1% thioflavin solution, diluted in the 80% of ethanol, for 15 min. The sections were then washed with ethanol and stained with DAPI. The green fluorescence of thioflavin S stained plaques were visualized with a Zeiss M2 upright microscope. The number of Aβ plaques were counted within the same size field of view in μm^2^ under the microscope. Quantitative densitometry analysis was performed using Image J (NIH, Version 1.43u) to semi-quantitatively compare the density of Aβ plaques in the brain.

### Primary cortical neuron culture

Primary cortical neuron culture was performed as previously described, with minor modifications ^29^. Briefly, E15 embryonic mouse cerebral cortex was treated with papain (P3125-10MG, sigma) dissolved in Neurobasal^®^ Media (Gibco, USA) for 30 min at 37 °C to dissociate cortices into single cells. Cells were then resuspended in neurobasal media containing trypsin inhibitor. Tissues not sufficiently digested were removed after a brief centrifugation. Fully separated cortical neurons in the supernatant were collected and plated at 5 × 10^4^ cells/mL in 6-well (4 mL/well), 24-well (1 mL/well), or 96-well (100 μL/well) plates (Corning, NY). Neurons were cultured at 37 °C in a humidified atmosphere of 5% CO_2_ for 7 - 8 days before use.

### Ratiometric Measurement of [Ca^2+^]i using Fura-2

Ratiometric calcium imaging was performed on neurons to detect intracellular calcium level as we have previously described ^29^. Briefly, cortical neurons on coverslips in 24 well plates were treated with Fura-2-AM fluorescence dye (molecular probes^®^, Life Technologies, USA) in a final concentration of 5 μL/mL, and incubated at 37°C with 5% CO_2_ for 20 min. After rinsing twice with 37 °C PSS Mg^2+^-free buffer, the coverslips were transferred into a new 35 mm dish containing 3 mL PSS Mg^2+^-free buffer. Fura-2 fluorescence was measured at 510 nm emission with 340/380 nm dual excitation selected by a DG-5 system (Sutter Instrument Company, Novato, CA) and imaged with fluorescence microscope (Leica DMI4000B, Germany). The intracellular calcium ([Ca^2+^]i) concentration was expressed as ratios from fluorescence intensity between the two excitation wavelengths of R340/380 of Fura-2. Mg^2+^-free buffer containing 45 mM KCl was applied to neurons to induce [Ca^2+^]i in order to confirm neuronal membrane potential and viability following all treatments. All measurements were independently repeated for at least 3 times. For each experiment, at least 30 neurons were selected for analysis.

### Thioflavin T (ThT) fluorescence assay

Thioflavin T (ThT) fluorescence is used regularly to quantify the formation and inhibition of amyloid fibrils in the presence of anti-amyloidogenic compounds such as polyphenols. Thioflavin T (ThT) assay was performed as previously described with minor modifications ^30^. Briefly, Aβ_1-42_ powder was dissolved in 1mL 1,1,1,3,3,3-Hexafluoro-2-propanol (HFIP), sonicated for 5 min to completely dissolve the Aβ_1-42_, followed by incubation for 1 h at room temperature, and aliquoted into 1.5 mL microcentrifuge tubes. HFIP in each tube was completely evaporated over 3 h in a fume hood, followed by evaporation for 1 h in a SpeedVac concentrator (CentriVap^®^ Concentrator, Labconco, USA) at the room temperature. To produce aggregated Aβ_1-42_, the HFIP-treated Aβ_1-42_ peptides was re-dissolved in 10 mM NaOH and sonicated in cold water bath for 30 min, diluted to a final concentration of 25 μM in PBS (pH 7.4). The Aβ_1-42_ solution was incubated for 3 d at 37 °C to form aggregated Aβ_1-42_.

Aβ_1-42_ fibrils at 20 μM concentration were mixed with different concentrations of GA. The mixture was added into a 96-well plate at 200 μL/well, followed by incubating with 10 μM ThT for 15 min at 37 °C before recording. Fluorescence from ThT was detected using a Perkin-Elmer Luminescence spectrometer (EnSpire™ Multilabel Reader, Perkin-Elmer, Singapore). Excitation and emission wavelengths for fluorescence spectroscopy were at 450 nm and 485 nm, respectively.

### Dynamic light scattering (DLS) assay

DLS measurements were performed using a Zetasizer Nano S (Malvern Instruments, Worcestershire, UK) according to the method described in ^22^. For *in situ* measurements, autocorrelation functions of scattered light were collected using acquisition times of 3 min and converted into particle-size distributions using the “narrow modes” or “general purpose” algorithms provided with the Zetasizer Nano S. Changes in scattering intensity were derived from the count rates of the avalanche photodiode photon detector.

### DLS measurements

Aggregation state of Aβ_1-42_ with the presence of GA was further determined using DLS technique. The hydrodynamic radius (Rh) measurements were taken at 90° angle and 637 nm on Brookhaven BI-200SM. All samples (Aβ_1-42_ fibrils at 20 μM and GA at 40 μM) were filtered through a 0.22 μM pore sized micro filter (Millex-GP, SLGP033RB) and placed directly into a quartz cuvette. Data collection was done at a physiological temperature (37°C). The mean Rh and the polydispersity (Pd) were calculated from the autocorrelation analysis of light intensity data (>40 acquisitions) based on the diffusion coefficients, which can be converted by the Stokes-Einstein equation.

### Atomic force microscopy (AFM)

Aβ_1-42_ solution at 10 μL volume was loaded on freshly cleaved, unmodified AFM grade mica surface (Electron Microscopy Sciences, cat. No. 71856-01-10, USA). Samples were incubated with mica for 10 min and rinsed with 5 drops of deionized water to remove buffer salts and unbound peptides. The mica surface was blow-dried under a gentle stream of nitrogen. The AFM experiments were conducted in tapping mode in air on an MFP-3D Infinity AFM (Asylum Research, Oxford Instruments) at room temperature.

### Animal model of AD

All animal experiments were approved by the Animal Ethics Committee of the Southern University of Science and Technology (Shenzhen, China). APPswe/PS1dE9 (referred to as APP/PS1) transgenic mice were gifts from Institute of Neuroscience at Shanghai (Chinese Academy of Sciences). These mice with a C57BL/6 background express chimeric amyloid precursor protein (APPswe) encoding the Swedish mutations at amino acids 595/596 and an exon-9-deleted human PS1 (PS1dE9) controlled by independent mouse prion protein promoter elements ^31^. Both of the genetic modifications are associated with familial AD in humans ^32^. Mice were maintained in a SPF II animal facility with controlled environment (temperature: 22 ± 1°C; humidity: 50 ± 10 %, and a 12 h light/dark cycle). Mice were allowed food and water *ad libitum*. These mice were bred with C57BL/6 mice to establish a colony of APP/PS1 mice and wildtype littermates. Genotyping of mice was carried out using tail tip DNA following the standard DNA extraction and PCR protocol.

### Gallic acid administration

We used APP/PS1 and their wildtype littermates to examine whether GA could ameliorate AD pathology and cognitive deficits *in vivo*. GA treatment was given to both the 4 month-old group and the 9-month-old groups. GA was dissolved in sterile water at the concentration of 3 mg/mL, and was given to mice daily at 30 mg/kg through gavage as previously described ^25^. Sterile water was given to mice serving as a vehicle-treated control group. All mice were weighted every week in the morning. After GA treatment, mice were subjected to a suit of behavioral tests.

### Mouse behavioral tests

**1. Open field test:** After the last GA administration, mice spontaneous activity in the open field were tested using a clear plastic cube box (41× 41× 38 cm) under a camera, the center region was defined as a 20 cm × 20 cm virtual area. Each mouse was given 10 min to move freely in the box. All traveled distances and the distance trace center was recorded and analyzed using Smart V3.0, Panlab software. The rearing number, representing the number of times mouse lifting the forepaws to explore upwards, was recorded by the experimenter.

**2. Morris water maze (MWM):** Morris water maze was performed to determine the spatial learning ability and reference memory. MWM was performed in a blue water pool (120 cm in diameter with 2/3 of transparent water) containing a circular bright blue platform (14 cm in diameter and submerged 1.5 cm beneath the water surface).

*Spatial acquisition test:* The spatial acquisition learning ability training consisted of five consecutive days of training with every training day comprising of four trials with 15 min inter-trial interval. The entry points randomly selected each time from the different designated locations. Once mouse successfully found the platform within 60 s, it was placed into a cage under a warming lamp. Otherwise, mouse was gently and manually guided to the platform and allowed to remain there for at least 20 s. The traveled distance to platform was recorded in each day to assess the spatial learning ability.

*Probe trial test:* On day 6, a probe trial test was performed. The hidden platform was removed from the pool and the mouse was placed in the quadrant diagonally opposite the target quadrant and allowed to swim for 60 s. The time spent in the target quadrant and mean distance to target were recorded and analyzed as a measure of spatial memory retention (or the reference memory) ^33^. Mice were monitored by a camera and its trajectory was analyzed using the Smart V3.0, Panlab software.

**3. Y-maze spontaneous alternations:** To assess spatial working memory of the mice, spontaneous alternation behavior was assessed in a Y maze after the MWM. The Y maze used in this test was composed of three identical acrylic light blue arms (31cm long, 17 cm high and 9.5 cm wide). During each trial, mouse was placed at the maze center and allowed to explore freely through the maze for 5 min. The sequence of arm entries was recorded by a monitor right above the apparatus and analyzed using the Smart V3.0, Panlab software. Alternation percentage (%) was calculated as a proportion of arm choices differing from the previous two choices ^34^.

**4. Novel object recognition test:** The novel object recognition test was carried out to assess the short-term recognition memory after the Y maze test. The task was adapted to the experimental setting as described previously ^35^. Each trial of a mouse consisted of a familiarization session and a test session. In the familiarization session, each mouse was individually placed into an open field testing box, mouse was allowed to explore freely for 10 min in the box with two identical objects on the symmetrical diagonal position. One hour after the familiarization, one familiarized object in the box was replaced with a novel object having different brightness, shape and texture. In the test session, mouse was put back into the box again and given 5 min to explore the two different objects. The activity of the mice was recorded using Noldus video tracking system (Ethovision). Duration of time was recorded to represent basic tendency of the mouse to sniff and explore novelty objects when the mouse nose point was around the object (distance between the nose and the object was less than 2.5 cm). Discrimination index was defined as the ratio of exploration duration around the novel object over duration around both the objects in the test session.

In order to minimize the difference made by experimenter handling, throughout all behavioral experiments, mice were handled by the same one experimenter.

### Electrophysiological recordings of Long term potentiation (LTP)

The method used to obtain LTP was essentially based on a previously described protocol ^29,36^. The transverse hippocampal slices (400 μm) were prepared using a Vibrotome slicer (VT 1000S; Leica) in ice-cold artificial cerebral spinal fluid (aCSF). The composition of aCSF was (in mM): 120 NaCl, 2.5 KCl, 1.2 NaH_2_PO_4_, 2.0 CaCl_2_, 2.0 MgSO_4_, 26 NaHCO_3_, and 10 glucose, saturated with 95% O2 / 5% CO2 (v/v). After incubating for 30 min at 34 °C and 2 - 8 h at the room temperature (25 ± 1 °C), the hippocampal slice was kept submerged in aCSF (3 mL/min; 32-34 °C) with a nylon mesh. The same aCSF that injected into the glass pipettes (1-5 MΩ) was used to record the field potential, field excitatory postsynaptic potentials (fEPSPs) evoked in the CA1 stratum radiatum by stimulating Schaffer collaterals (SC) with a two-concentrically bipolar stimulating electrode. The evoked responses were recorded in current-clamp mode by the Axon MultiClamp 700B (Molecular Devices) amplifier. Stimulation intensity was adjusted to evoke fEPSP amplitudes that were 25% of maximal response. LTP was induced by applying one or four trains of high-frequency stimulation (HFS: 50 pulses in 100 Hz, and an inter-train interval of 10 s, stimulation at test stimulus intensity).

### GA Binding Docking Modeling

All GA derivatives were purchased from Sigma Aldrich. GA derivatives binding and interaction with Aβ_1-42_ were examined using Molecular Operating Environment (MOE) 2014.09 software at the atomic level ^37^. Crystal structure of fibrillar Aβ_1-42_ (PDB: 2MXU) was downloaded from PDB website, and was structure-prepared using Protonate 3D program in MOE. Induced fit was selected as the docking protocol, and Amber12: EHT was selected as the forcefield. The other settings were selected as default options. The docking results were scored based on London dG scoring function and the top one was selected for binding analysis.

### Statistical analysis

All data were represented as mean ± SEM. One-way ANOVA plus LSD *post hog* analysis were employed to analyze the data obtained from body weight measure, probe trial test of MWM, OFT, Y maze, NORT, quantification of Aβ plaques. A repeated measure ANOVA was applied to the analysis of traveled distance to platform in the spatial acquisition test of MWM in order to detect significant differences between groups at different days. For this test, LSD *post hoc* test was performed. For the LTP recordings, neuronal viability and ratiometric measurement of [Ca2+]i data analyses, one-way ANOVA plus Tukey’s *post hog* analysis was used. All analyses were performed using SPSS 22.0 software. A P value of < 0.05 was taken to indicate statistical significance.

## Results

### 1. GA alleviates cognitive impairments of early stage AD mouse

To determine the possible beneficial effect of GA in the early stage of AD development, 4-month-old APP/PS1 transgenic mice were give GA through gavage for 30 d at 30 mg/kg/d. The drug dosage and route of delivery were chosen based on a previous report of similar animal studies ^25^ and our *in vitro* studies. Mice were randomly assigned into four groups (n = 6 – 9 per group): WT group, wildtype mice; WT+GA group, wildtype mice treated with GA; APP/PS1 group, AD transgenic mice; and APP/PS1+GA group, AD transgenic mice treated with GA (Fig 1). All groups of mice were monitored for physiological changes during the period of oral administration of GA: (1) Mice were weighted every week to confirm no significant changes in body weight (Supplemental Fig. S1A). (2) The open field test (OFT) was performed at the end of the last GA oral administration to monitor the spontaneous activity of GA-treated mice. All four groups showed no significant differences (Supplemental Fig. S1B-D). Cerebral blood flow of all groups was measured using Moor FLPI-2 (full-field laser perfusion imager) to exclude the possibility that GA may affect cerebral blood flow (Supplemental Fig. S3). These experiments indicated that GA treatment did not have a major negative impact on mouse normal brain physiology.

**Fig 1:**
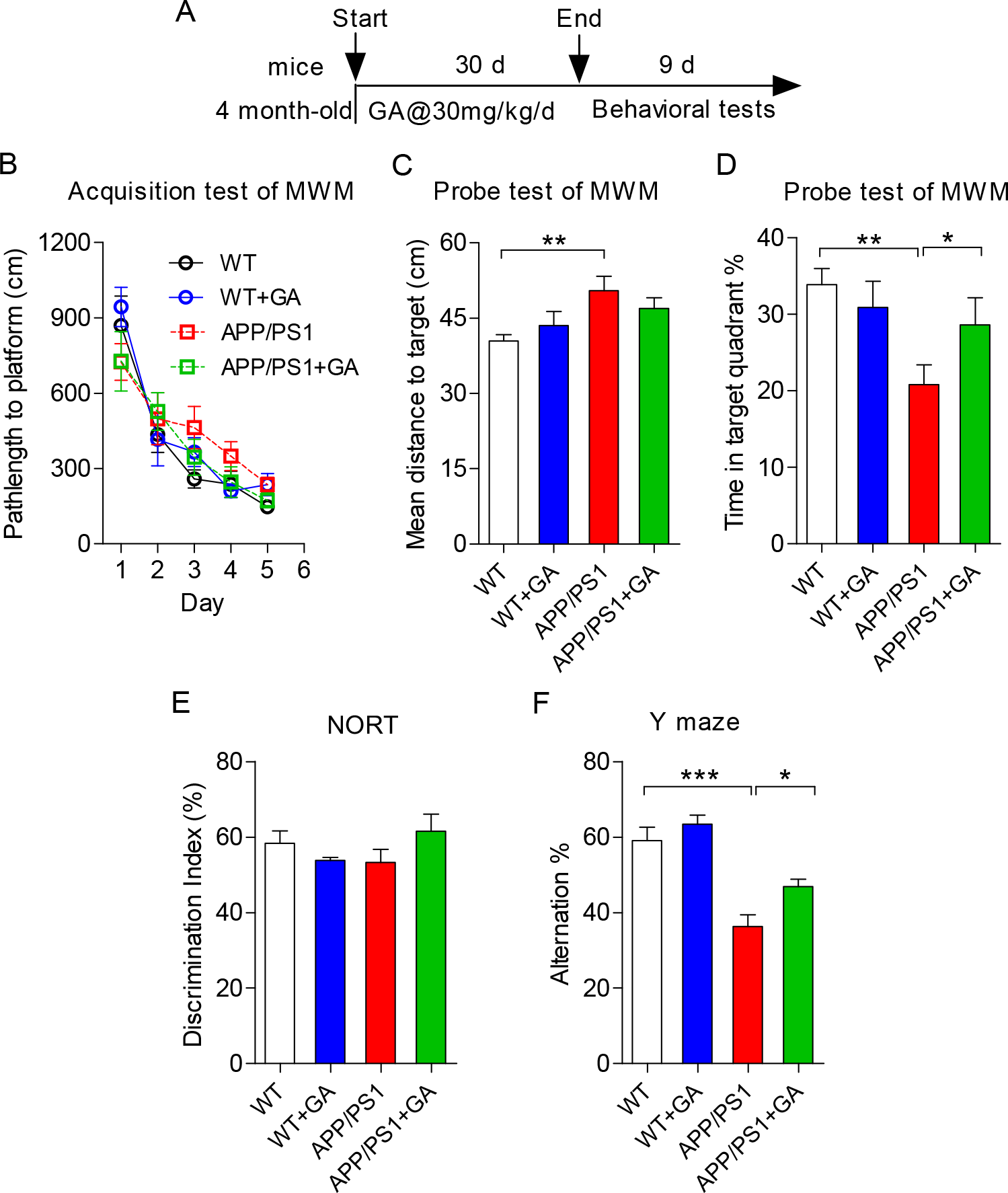
GA alleviated the mild cognitive impairment in APP/PS1 mice. (A) Experimental design: Starting at 4 months of age, WT littermates and APP/PS1 transgenic mice were administrated with GA at 30 mg/kg/day for 1 month, followed by a battery of behavioral tests lasting for 9 days. Behavioral tests included MWM (B-D), NORT (E), Y-maze (F). (B) shows the acquisition test of the MWM. Mean path-length to the platform was calculated for each day at the spatial acquisition stage of MWM of all four groups of mice. (C) Mean distance to the target, and (D) Percentage of time spent in the target quadrant during the spatial exploring stage of MWM are shown. (E) The percentage of exploration time spent on the novel object in the test session (NORT), expressed as a discrimination index, is shown in panel E. (F) The percentage of spontaneous alternation in Y maze was recorded and calculated shown in panel F. Data represent mean ± SEM. Error bars indicate SEM (n = 6-10 mice in each group). * p < 0.05, * * p < 0.01, * * * p < 0.001.

Subsequently, a battery of behavioral tests was carried out on these mice as depicted in Fig 1A. Behavioral tests included Morris water maze test (MWM), novel object recognition test (NORT) and the Y maze test. As shown in Fig 1B, the spatial acquisition profiles of all 4 groups of mice were similar with a gradual shortening of path length to platform measurements over the 5 d spatial acquisition training period. These results demonstrated that the spatial learning abilities of APP/PS1 transgenic mice were comparable to those of the WT littermates. But, in the subsequent day 6 of spatial probe trial test, the APP/PS1 transgenic mice group exhibited mild deficit in memory retention as shown by the appearance of more random location searches, a longer mean distance of path length to target (p < 0.01, Fig 1C), and a lesser amount of time spent in the target quadrant (p< 0.01, Fig 1D) compared to the WT littermate mice (Fig 1C, D). By contrast, GA-treated group of APP/PS1 mice (APP/PS1+GA group) spent significantly more amount of time in the target quadrant (p < 0.05, Fig 1D) compared with those of APP/PS1 group. These data demonstrated that GA improved the spatial reference memory deficit in the young AD mice.

In addition, NORT was performed which showed no significant differences amongst all groups in the discrimination index, indicating clear preferences for the exploration of novel objects during the test session (Fig 1E). In contrast, Y-maze test, which evaluates spatial working memory, showed that the APP/PS1 group exhibited a significantly lower alternation behavior than the WT group (p < 0.001, Fig 1F), confirming deficits in remembering which arm was explored. This alternation behavior deficit was significantly ameliorated in the APP/PS1+GA group (p < 0.05, Fig. 1F), demonstrating that GA was effective in enhancing spatial working memory during the early stage of AD development.

### 2. GA alleviates cognitive impairments of late stage AD mouse

To determine whether GA can also reduce the more severe cognitive deficits of AD mice at a relatively late stage, the 9-month-old APP/PS1 transgenic mouse was given GA orally through gavage for 30 d at 30 mg/kg/d. The treatment schema is shown in Fig 2A, which was identical to that used for the 4-month-old mice. Again, no drastic changes in the body weight and in both GA-treated WT and APP/PS1 mice as shown in the Supplemental Fig S2A (body weight measurements). Based on OFT data, the 9-month-old APP/PS1 mice showed more exploratory activities, representing increased anxiety compared to the WT littermates (Supplemental Fig S2B-D). Interestingly, GA treatment reduced this behavior in APP/PS1 mice as show in Supplemental Fig S2 B-D.

**Fig 2:**
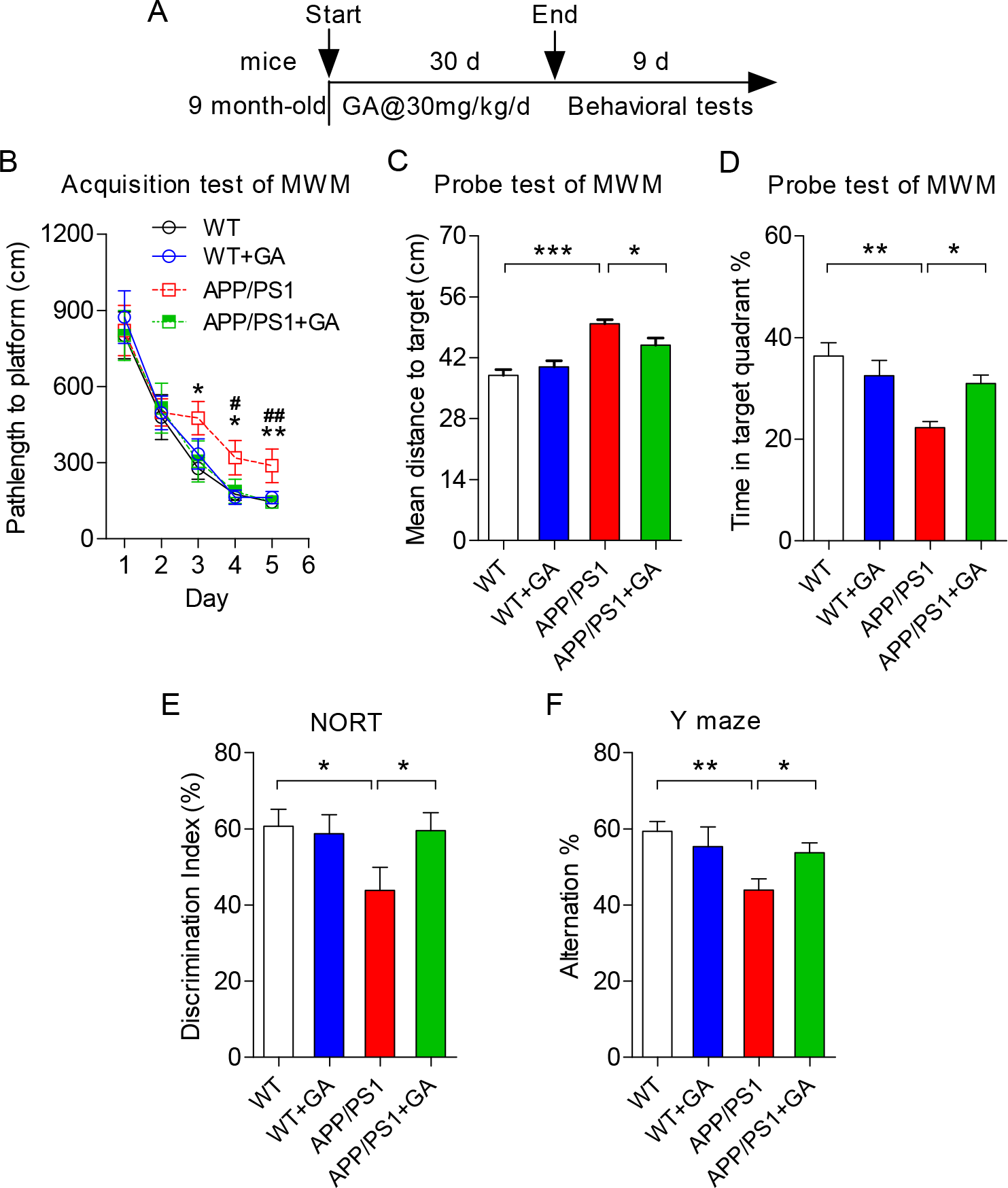
GA alleviated the cognitive deficits of late stage APP/PS1 mice. (A) Experimental design: Starting at 9 months of age, WT littermates and APP/PS1 transgenic mice were administrated with GA at 30 mg/kg/day for 1 month, followed by a battery of behavioral tests lasting for 9 days. Behavioral tests included MWM (B-D), NORT (E), Y-maze (F). (B) shows the acquisition test of the MWM. Mean path-length to the platform was calculated for each day at the spatial acquisition stage of MWM of all four groups of mice. (C) Mean distance to the target, and (D) Percentage of time spent in the target quadrant during the spatial exploring stage of MWM are shown. (E) The percentage of exploration time spent on the novel object in the test session (NORT), expressed as a discrimination index, is shown in panel E. (F) The percentage of spontaneous alternation in Y maze was recorded and calculated shown in panel F. Data represent mean ± SEM. Error bars indicate SEM (n = 6-10 mice in each group). * p < 0.05, * * p < 0.01, * * * p < 0.001.

The 9-month-old APP/PS1 mice exhibited significant cognitive impairments before GA treatment (Fig 2B-F). The spatial learning ability of APP/PS1 mice was significantly impaired based on the spatial acquisition trials of the MWM test (Fig 2B, p < 0.05 on the 3^rd^ and 4^th^ d, p < 0.01 on the 5^th^ d). Poor memory retention with a longer mean distance of path length to target (Fig 2C, p < 0.001) and a lesser time spent in target quadrant (Fig 2D, p < 0.01) occurred in APP/PS1 mice during the probe trial tests when compared with those of the WT group. Remarkably, GA treatment of APP/PS1 transgenic mice significantly enhanced the acquisition of spatial memory compared to APP/PS1 group (Fig 2B, # indicates p < 0.05; ## indicates p < 0.01). GA-treated APP/PS1 mice also showed better memory retention in the probe trials than the untreated mice (APP/PS1 group, Fig 2C and D). These data indicated that GA can mitigate the spatial learning and reference memory deficits in late stage of AD development.

The discrimination index of the APP/PS1 mice in the NORT test showed significant reduction compared with that of the WT mice (Fig 2E, p < 0.05). After GA administration, the percentage of recognition index was significantly enhanced in the GA-treated APP/PS1 group (Fig 2E, p < 0.05). Moreover, the alternation percentage of APP/PS1 group was significant reduced compared with WT group and APP/PS1+GA group in the Y-maze test (Fig 2F, p < 0.01 with WT *vs*. APP/PS1; p < 0.05 in APP/PS1 *vs*. APP/PS1+GA). These findings suggested that GA was effective in enhancing both the short-term recognition memory and the spatial working memory in late stage AD mice.

### 3. GA improves hippocampal LTP in the late stage AD mice

LTP is widely considered as one of the major cellular mechanisms that underlies learning and memory. To understand GA’s mechanism in enhancing spatial learning and memory, electrophysiological recordings of LTP in response to GA treatment were performed. High frequency tetanic stimulation of the Schaffer collaterals projected to the CA1 pyramidal neurons in acute hippocampal slices established the baseline synaptic responses of 10-month-old WT and APP/PS1 littermate mice. The input/output (I/O) curves of field excitatory postsynaptic potentials (fEPSP) slope with series of increasing stimulation intensities were shown in Fig 3A. Comparing the basal synaptic transmission between the WT littermates with APP/PS1 mice showed no statistically significant difference (Fig 3A).

**Fig 3:**
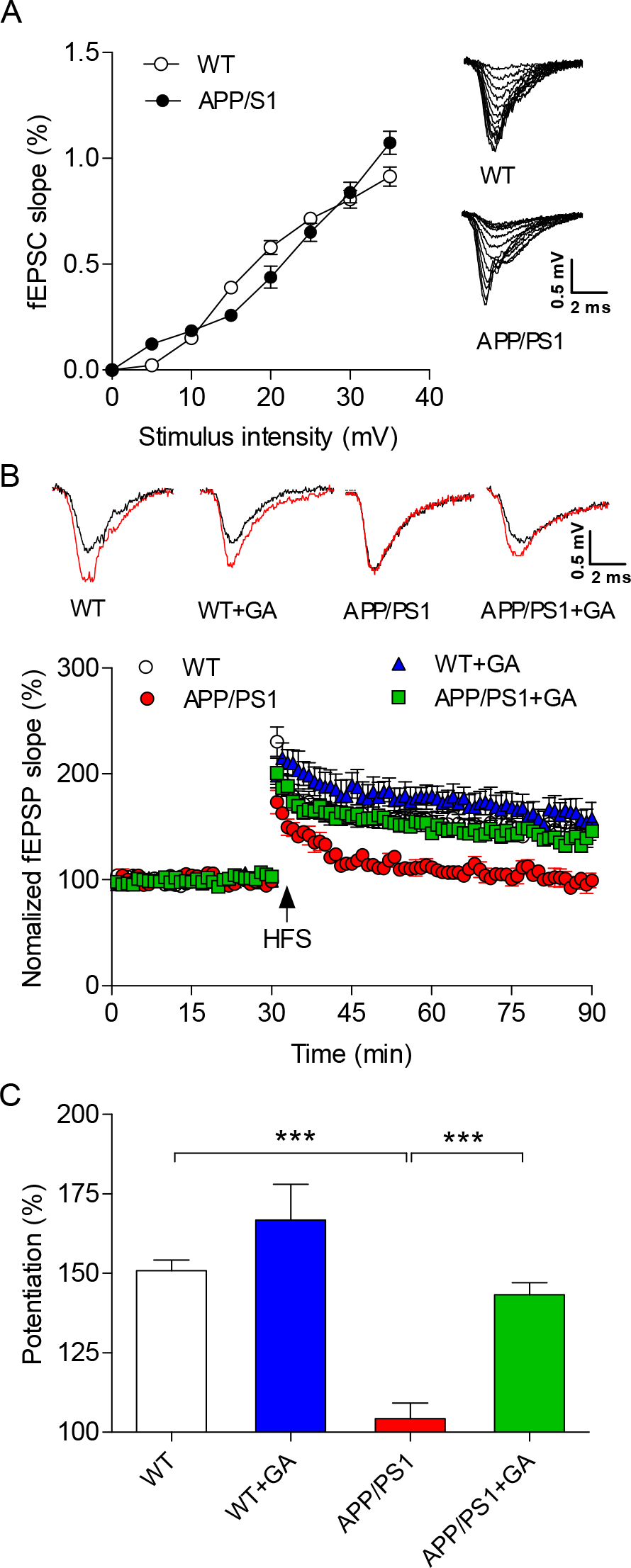
GA treatment significantly improved the synaptic strength of 10-month-old APP/PS1 mice. The LTP was generated by high frequency tetanic stimulation (HFS) of the Schaffer collaterals projected to the CA1 pyramidal neurons on acute hippocampal slices from the 10-month-old APP/PS1 mice. (A) Input/output [IO] curves of fEPSP (mV/ms) versus stimulation intensity (mV) were taken from the wild-type littermate and APP/PS1 mice. The changes fEPSPs slopes of the four groups over the time course are shown in panel (B). The first 20 min of evoked responses were normalized and used as the baseline responses of LTP. (C) The magnitude of LTP was determined according to the responses between 45 and 60 min after the HFS. Data represent mean ± SEM. Error bars indicate SEM (n = 5 -6 mice in each group). * * * p < 0.001.

However, LTP generated in the 10-month-old APP/PS1 group of mice was significantly reduced compared to the WT littermate group of mice (Fig 3B and C; p < 0.001 with WT potentiation at 150.86 ± 3.33% vs. APP/PS1 potentiation at 104.29 ± 4.81%), confirming the development of severe deficits in synaptic strength associated with learning and memory in the 10-month-old APP/PS1 mice. Remarkably, APP/PS1+GA group significantly augmented the normalized slope of fEPSP compared with that of the APP/PS1 mouse group (Fig 3B and C; p < 0.001 with APP/PS1+GA group potentiation at 143.23 ± 3.75% vs. APP/PS1 potentiation at 104.29 ± 4.81%), suggesting GA treatment led to the enhancement of synaptic strength in APP/PS1 mice.

### 4. Gallic acid treatment reduces the Aβ plaque size in the APP/PS1 mice brain

To determine whether GA reduces Aβ load in the treated mouse brains, mouse brain tissue sections were stained with thioflavin S (ThS) which selectively labels Aβ plaques (Fig 4A). First, APP/PS1 mice showed a significantly increased presence of Aβ plaques in the CA1, DG and cerebral cortex regions (Fig 4A). There were no Aβ plaques found in the CA3 region (not shown). After one month of GA oral administration to the 9-month-old APP/PS1 mice (APP/PS1+GA group), the number of plaques per mm^2^ area in the hippocampal CA1 and DG regions remained unchanged compared to that of the GA-treated APP/PS1 mice (Fig 4B, C), except the number of Aβ plaques in the cerebral cortex, which showed significant reduction in the GA treated APP/PS1 mice group (Fig 4D). Interestingly, the average size of Aβ plaques was significantly reduced in APP/PS1+GA group compared with those of the APP/PS1 group in the CA1, DG and cerebral cortex regions (Fig 4E-G). These findings suggest that GA potentially prevented the formation of Aβ aggregation and plaque formation.

**Fig 4:**
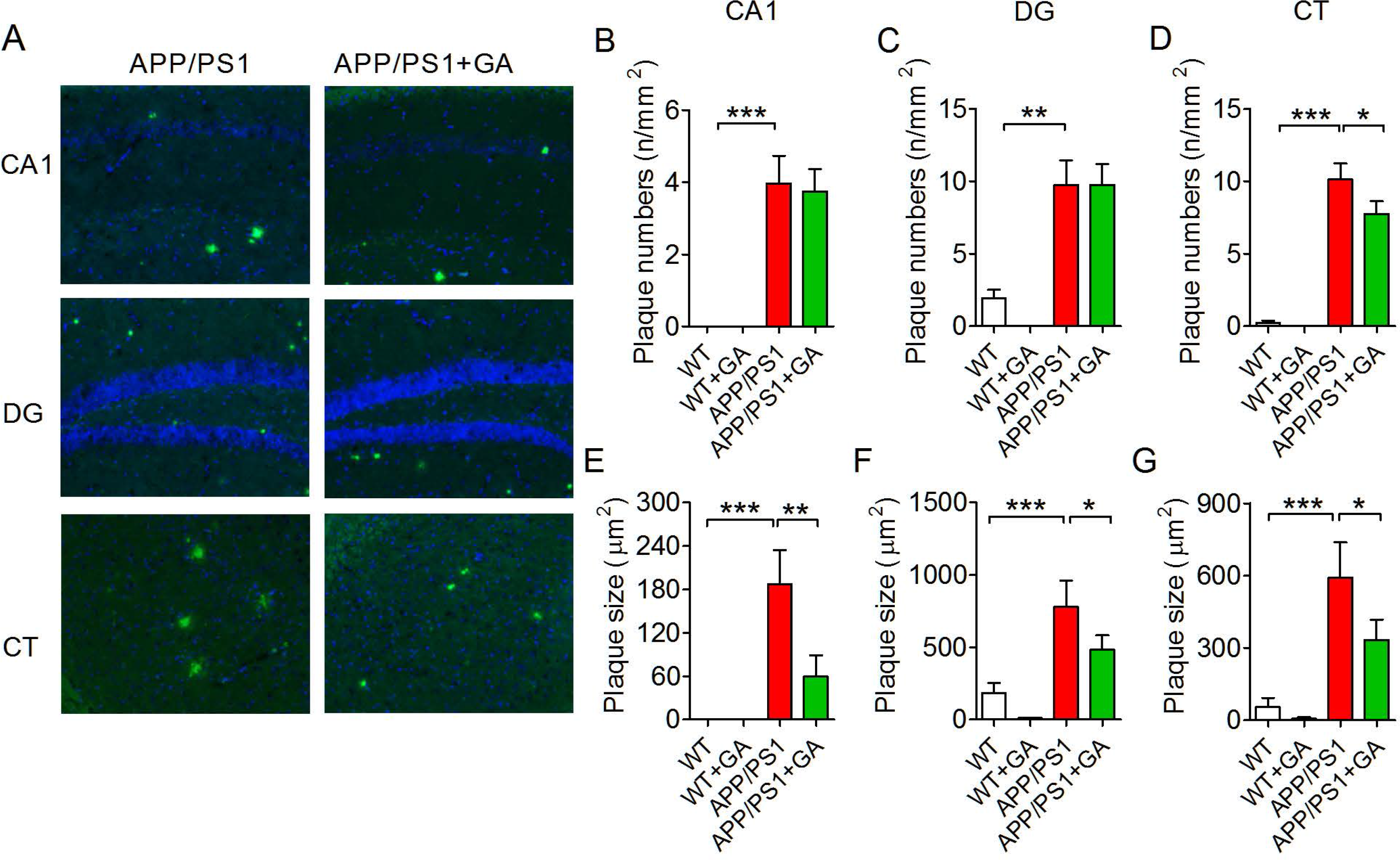
GA reduced the Aβ_1-42_ plaque size in hippocampus and cortex of ten-month-old APP/PS1 mice. (A) Representative images from the CA1 subfield, dentate gyrus (DG) and cortex (CT) of APP/PS1 and GA-treated APP/PS1 mice, which were stained using ThS to show Aβ plaques. (B-D) Aβ plaque numbers and (E-G) sizes in different regions were measured and quantitatively analyzed using Image J. Data represent mean ± SEM with n =3 in each group. * P < 0.05, * * P < 0.01, * * * P < 0.001.

### 5. Gallic acid decreases Aβ1-42 aggregation and Aβ_1-42_-induced neuronal death in vitro

To further investigate the possibility that GA can reduce Aβ aggregation, the following four *in vitro* experiments were performed. First, thioflavin T (ThT) fluorescence densitometry assay was used to quantify the formation of Aβ fibrils in the presence of GA. At the ratio of 1:2 (Aβ:GA) and to a lesser extend at 1:1 (Aβ:GA), GA significantly inhibited the formation of Aβ fibrils over time in solution (Fig 5A). Second, dynamic light scattering (DLS) was used to determine the size distribution profile of Aβ particles in solution suspension. Adding GA to the aggregated Aβ_1-42_ fibrils (Fig 5B) for 2 h clearly reduced the Aβ_1-42_ fibril particle size from predominantly 100 nm fibrils (blue colored bars) to about 60 nm sized particles (red colored bars). Third, the atomic force microscopy (AFM) technique was used to show that adding GA to the aggregated Aβ_1-42_ fibrils (Fig 5C) or co-incubating with Aβ_1-42_ clearly reduced the Aβ_1-42_ fibril particle size distribution profile. Fourth, indeed, the neurotoxicity of oligomeric Aβ_1-42_ mixtures, but not the fibrillar Aβ_1-42_, was significantly reduced in *in vitro* cultured cortical neurons by GA (Fig 5D). Neuronal death of different kinds of Aβ_1-42_ mixtures were shown on each corresponding AFM pictures in Fig 5C and the corresponding viability shown in Fig 5D. Additionally, cultured cortical neurons preincubated with GA for 15 min before the addition of oligomeric forms Aβ_1-42_ also significantly protected against Aβ_1-42_ toxicity (Fig 5E). At this condition, Aβ_1-42_ at 2.5 μM concentration killed 50% of cultured cortical neurons over 24 h period (Fig 5E). In contrast, GA at 1 and 5 μM concentrations provided significant neuroprotection (p < 0.001). Glutamate-induced neurotoxicity and MK801 protection against glutamate toxicity were used as internal controls (Fig 5E).

**Fig 5:**
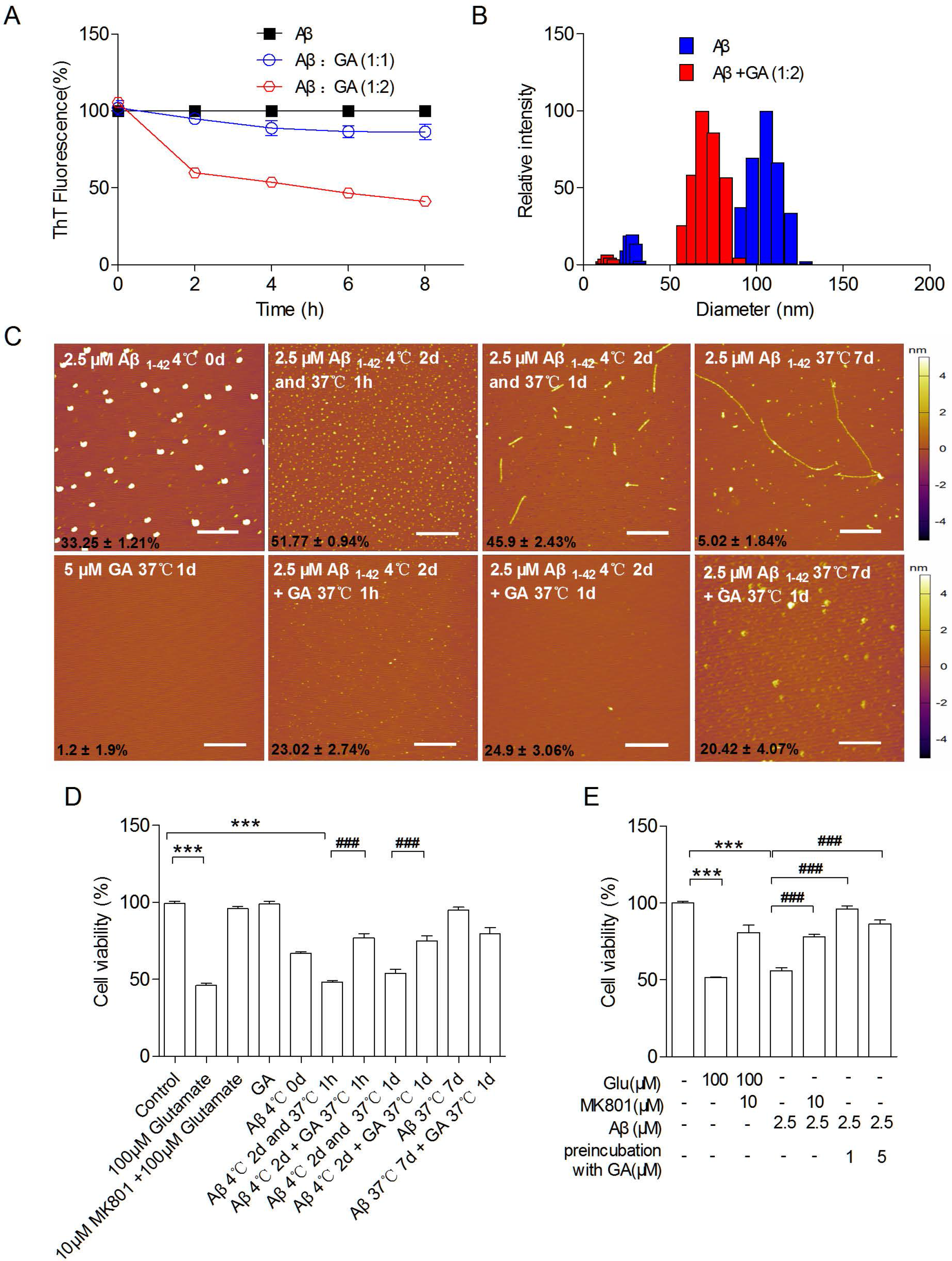
GA reduced fibrillar Aβ_1-42_ aggregation and toxicity to neurons. GA at the specified concentrations were mixed with the aggregated Aβ_1-42_ before the ThT fluorescence assay (A). (B) DLS assay was also performed to show the reduction of Aβ_1-42_ aggregate particle sizes when mixed with GA at the specified concentrations (B). (C) Top panel show the AFM images of fibrillar Aβ_1-42_ development. The bottom panel shows when GA effectively disrupted the formation of Aβ_1-42_ fibrils. Numbers at the left bottom corner refer to neuronal mortality rate caused by the mixture. (D and E) Neuronal viability was determined using CCK8 Viability Assay Kit. Pre-incubation with GA reduced the cytotoxicity caused by Aβ_1-42_. All data sets were n = 3. Values represented as mean ± SEM. Scale bars = 30 μm.

Taken together, these data clearly demonstrated that GA can effectively reduce Aβ_1-42_ fibril aggregation formation and neurotoxicity, which serves to explain the mechanism of GA’s effect in neuroprotection.

### 6. Oligomeric Aβ_1-42_ dose-dependently increases [(Ca^2+^)i] in cultured cortical neurons

It is known that oligomeric Aβ_1-42_ can induce i[(Ca^2+^)i] influx in cultured cortical neurons resulting in neuronal death ^12^. Therefore, the following experiments were designed to establish the dose-response relationship of [(Ca^2^+)i] changes in response to oligomeric Aβ_1-42_, and to determine whether GA treatment can reduce oligomeric Aβ_1-42_ toxicity to neurons through inhibiting [(Ca^2+^)i] changes.

First, Aβ_1-42_ peptide was polymerized *in vitro* as described in the Methods section to form oligomeric Aβ_1-42_. When added to the cultured neurons, ratiometric calcium imaging technique was used to show a dose-dependent increase in [(Ca^2^+)i] (Fig 6A). The dose-dependent relationship was determined in Fig 6B with the EC_50_ = 2.73 ± 1.047 μM. This experiment allowed the establishment of 2.5 μM Aβ_1-42_ as a working concentration for the oligomeric Aβ_1-42_ toxicity studies.

**Fig 6:**
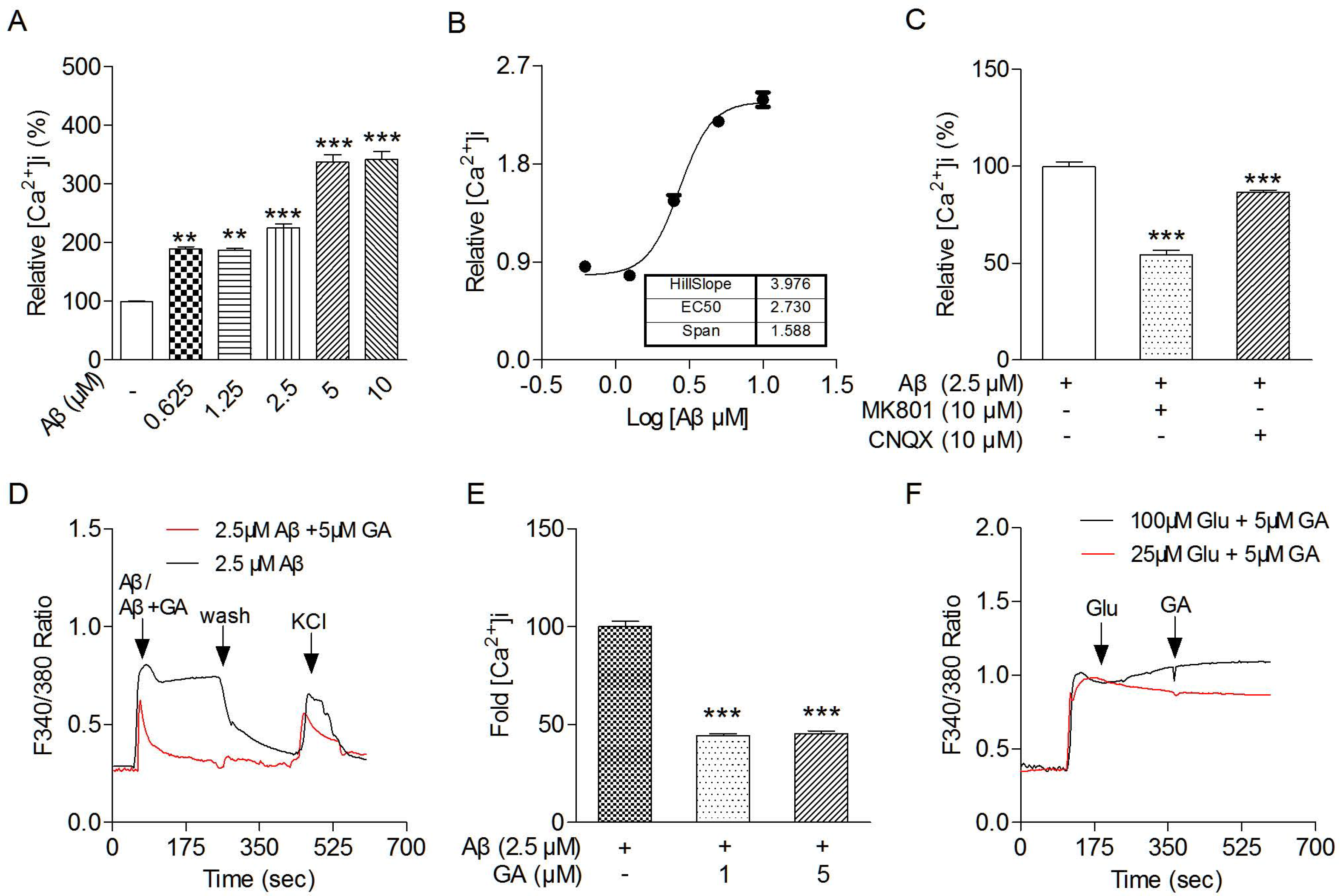
Aβ_1-42_ elicited increased intracellular calcium in primary cortical neurons. (A) Ratiometric calcium imaging measurement of the intracellular level of calcium ([Ca^2+^]i) in response to Aβ_1-42_ at doses ranging from 0.625 μM to 10 μM. The relative [Ca^2+^]i level of Aβ_1-42_ treatment was normalized to the baseline calcium level of non-treated cortical neurons. Data represents the average of measurements of at least 20 cells. (B) Concentration-response relationship for Aβ_1-42_-induced [Ca^2+^]i was determined and the concentration-response curve was fitted using the logistic equation. The EC_50_ was determined to be at 2.73 ± 1.047 μM and the slope factor of 3.976 ± 0.9103. (C) Cultured cortical neurons were treated with 2.5 μM of the oligomeric form of Aβ_1-42_. Fold reduction of [Ca^2+^]i in neurons treated with 10 μM MK801 or 10 μM CNQX in response to Aβ_1-42_-induced [Ca^2+^]i was measured. (D) Representative traces of [Ca^2+^]i in cortical neurons treated with oligomeric Aβ_1-42_ (2.5μM) (black colored line), or with pre-mixed GA with oligomeric Aβ at a ratio of 2:1 (5 μM : 2.5 μM). A brief wash with PSS removed the increase of Aβ_1-42_-induced [Ca^2+^]i in neurons. Addition of KCl showed a large increase in [Ca^2+^]i confirming that these neurons were functional cells. (E) Cultured cortical neurons were treated with oligomeric Aβ_1-42_ (2.5 μM) pre-mixed with GA at 37 °C for at least 15 min before measuring Aβ_1-42_-induced [Ca^2+^]i in neurons. (F) Cultured cortical neurons were treated with 25 μM or 100 μM glutamate followed by the addition of GA, which did not inhibit glutamate-induced calcium influx. Data represent mean ± SEM with n = 20 cells in each group. * * P < 0.01, * * * P < 0.001.

Second, inhibitors to NMDA receptors (MK801) and AMPA receptors (CNQX) were used to determine if they can block Aβ_1-42_ induced [(Ca^2+^)i] influx. As shown in Fig 6C, compared to the level of [(Ca^2+^)i] influx caused by 2.5 μM Aβ_1-42_, MK801 (10 μM) inhibited about 40% of that, while CNQX (10 μM) inhibited less than 20% of that, indicating that [(Ca^2+^)i] in neurons are through multiple channels in response to Aβ_1-42_. Furthermore, extracellular calcium is the main source for the increase of [(Ca^2+^)i] level seen in response to Aβ, as bathing neurons in calcium-free solutions showed no [(Ca^2+^)i] influx (Data not shown).

### 7. GA attenuated oligomeric Aβ_1-42_-induced [Ca^2+^]i in cultured cortical neurons and reduced neurotoxicity

To determine whether GA inhibits oligomeric Aβ-induced [(Ca^2^+)i] influx to neurons, cultured cortical neurons were treated with 2.5 μM oligomeric Aβ_1-42_ pre-mixed with GA (Fig 6D,E). Aβ_1-42_ used in this study was pre-polymerized and toxic to neurons as described in the Method section. From Fig 6D, it was clear that addition of oligomeric 2.5 μM Aβ_1-42_ elicited a fast and significant increase in [(Ca^2+^)i] influx to neurons, while GA with concentration ratios to Aβ at both 1:1 and 1:2 rapidly and significantly reduced [(Ca^2+^)i] influx into neurons. The inhibition by GA was about 50% of the [(Ca^2+^)i] influx level caused by Aβ_1-42_ without GA. These studies strongly suggest that GA was effective only at the level of reducing the polymerization of toxic Aβ_1-42_. Once polymerized, Aβ_1-42_ forms calcium conducting channels, GA would no longer be effective in blocking the action of polymerized Aβ_1-42_. Moreover, the fact that GA was not effective in blocking [(Ca^2+^)i] influx elicited by treatments with glutamate at 25 μM or 100 μM (Fig 6F), further supports the argument that the point of action for GA was not at the calcium channel level.

### 8. Molecular Modelling and Simulations of GA interaction with Aβ_1-42_

*Non-covalent interactions:* In order to provide a structural insight of GA disruption of Aβ_1-42_ aggregation, GA chemical derivatives were made which included 3-hydroxybenzoic acid, pyrogallol, 3,4-dihydroxybenzoic acid, 3,5-dihydroxybenzoic acid, and benzoic acid (Fig 7A). ThT fluorescence assay was used to determine whether these derivatives disrupt Aβ_1-42_ aggregation. As shown in Fig 7B and C, GA and pyrogallol significantly reduced Aβ aggregation-induced ThT fluorescence. In contrast, the rest of the GA derivatives was less effective (Fig 7C), suggesting that the three hydroxyl groups were essential in reducing Aβ_1-42_ aggregation.

**Fig 7:**
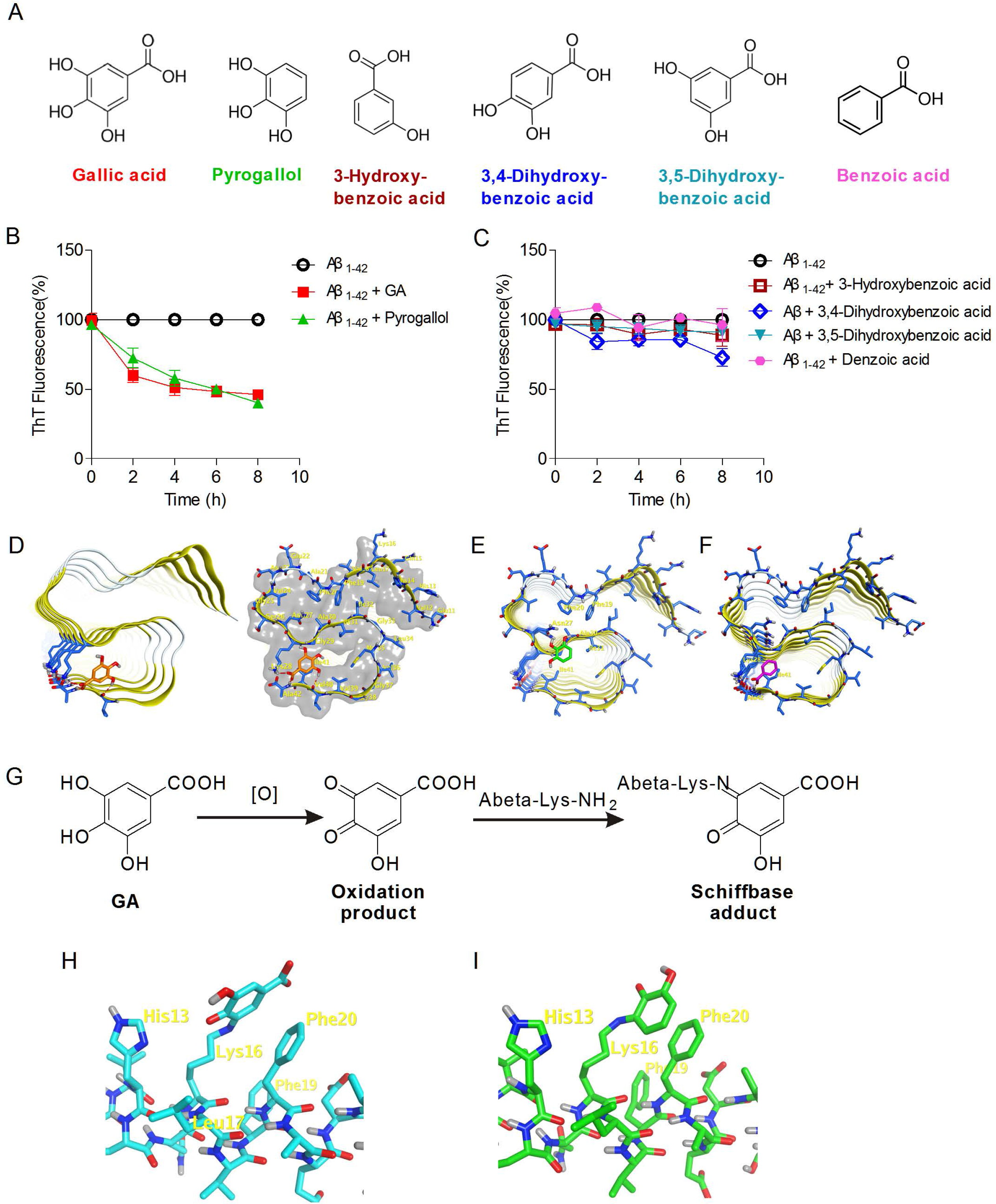
GA derivative structures (A), their effects in reducing ThT fluorescence intensity (B, C) and the predicted docking modes to fibrillar Aβ_1-42_ (D-I). GA derivative structures are shown in (A), where GA is in orange color; pyrogallol (benzene-1,2,3-triol) in green color; 3-hydroxybenzoic acid in brown color; 3,4-dihydroxybenzoic acid in blue color; 3,5-dihydroxybenzoic acid in paleturquoise color; benzoic acid in pink color. GA derivatives effect in reducing ThT fluorescence derived from Aβ_1-42_ aggregation (as 100%) is shown in (B) and (C) for clarity. Pyrogallol has similar potency to GA in reducing ThT fluorescence derived from Aβ_1-42_ aggregation (B), while all others were less effective in comparison (C). (D) Predicted binding modes of GA in the crystal structure of fibrillar Aβ_1-42_ shown as S-shaped triple-β-motif (PDB: 2MXU,) using the MOE software. GA is shown as orange colored stick; hydrogen bonding is represented as a red dash; Aβ_1-42_ is represented as yellow cartoon. Predicted noncovalent binding modes of pyrogallol (E, green sticks) and benzoic acid (F, pink sticks) in the aggregated structure of fibrillar Aβ_1-42_ (PDB: 2MXU) using the MOE software. (G) The oxidation product of GA derivatives and the proposed Schiff base adduct.Covalent docking of oxidized product of GA (H) and pyrogallol (I) with Aβ_1-42_ (PDB: 1IYT) through a Schiff-base formation between the carbonyl group of the oxidized product and ε-amino group of Lys16. Covalent docking was conducted with DOCKTITE program in the MOE software.

The crystal structure of fibrillar Aβ_1-42_ (PDB: 2MXU) is a S-shaped triple-β-motif, which is stabilized by a salt bridge between Lys28 and Ala42 residues. Docking studies revealed that the carboxylic acid group of GA forms hydrogen bond interactions with Lys28 and Ala42 residues, and these interactions were stabilized by hydrogen bonding with Val40 and hydrophobic interactions with Ile41 and Val39 residues (Fig 7D). The hydrogen bonding interactions between GA and Lys28 and Ala42 residues might break the salt bridge between Lys28 and Ala42, resulting in instability of fibrillar Aβ_1-42_, and ultimately lead to disaggregation of Aβ_1-42_ fibrils.

Although hydrogen bonding interactions were observed between benzoic acid and residues Lys28 and Ala42, no additional hydrogen bonding interaction was formed with other residues. Thus, interaction between benzoic acid and Lys28 and Ala42 may be not potent enough to break the salt bridge (Fig 7F). This suggests that the three hydroxyl groups play a role in stabilizing GA interaction with Aβ aggregation while the presence of the -COOH group is critical in destabilizing the Aβ_1-42_ fibrils.

Pyrogallol has additional binding sites on fibrillar Aβ_1-42_. Hydrogen bonding interactions were observed between pyrogallol and Asn27 and Ala30 residues (Fig 7E). In addition, Van der Waal interactions with Phe19, Phe20, Ile31 and Ile41 residues were also observed (Fig 7E). Indeed, the crystal structure of fibrillar Aβ_1-42_ revealed that the inter-chain hydrogen bonding between residue Asn27 is important for the aggregation and fibrils formation. Therefore, the stable hydrogen bonding interaction between pyrogallol and residue Asn27 may explain the disaggregation effect of pyrogallol on fibrillar Aβ_1-42_.

*Covalent interactions:* Another possible disaggregation mechanism of GA derivatives may be through covalent modification. As shown in Fig 7G, GA and pyrogallol contain a catechol moiety which is easily oxidized to quinone and is susceptible to form covalent interaction with one or more lysine residues on Aβ_1-42_ via Schiff-base formation. This covalent interaction will disaggregate the fibrillar Aβ_1-42_. To demonstrate this interaction, a covalent docking of GA quinone and pyrogallol quinone in monomers with Aβ_1-42_ was performed. As shown in Fig 7H and Fig 7I, both the oxidized product of GA and pyrogallol covalently interacted with Aβ_1-42_ through Schiff-base formation between the carbonyl group of the oxidized product and ε-amino group of Lys16. The van der Waals interactions between phenyl group of oxidized products and residues Phe20 and Phe19 contributed to the covalent modification. In addition, GA quinone forms an electrostatic interaction with His13. This may explain the lower activity of 3,4-dihydroxybenzoic acid than that of GA, as there is no hydroxyl group in the oxidized product of 3,4-dihydroxybenzoic acid, which means no hydrogen bonding interaction with His13 can be formed. These docking results also revealed that the oxidized product of GA and pyrogallol may be susceptible to form covalent interactions with Lys16, but not the Lys28 residue. We assumed that this may be due to the van der Waals interactions and electrostatic interaction with residues that are adjacent to Lys16.

## Discussion

In this work, we have showed what is to our knowledge the first detailed behavioral studies of GA’s beneficial effect on alleviating cognitive decline of APP/PS1 transgenic mouse, which have been hitherto unavailable. Series of mechanistic analysis showed that GA can disrupt Aβ_1-42_ aggregation thereby reducing neurotoxicity. Despite of the moderate resolution, molecular docking of interactions between GA and its derivatives with Aβ_1-42_ aggregates demonstrated that the three hydroxyl groups of GA are essential in stabilizing the interaction of GA with Aβ_1-42_ aggregation. Neurotoxicity in AD brain mainly comes from the oligomerized and aggregated forms of Aβ_1-42_. A drug with the capacity to disrupt Aβ_1-42_ oligomerization and aggregation represents a good candidate as it may prevent and treat AD. Indeed, one of the most interesting findings of the study was that GA not only significantly improved the spatial reference memory and spatial working memory during the early stage of AD development (4-month-old APP/PS1 mice), but can also significantly treat cognitive deficits in spatial learning, reference memory, short-term recognition and spatial working memories in the late stage AD mice (9-month-old). These studies indicated to us that GA can potentially both prevent and treat AD. Our data showed that the neuroprotective functions of GA were mostly through its ability to reduce Aβ_1-42_ aggregation from forming toxic oligomers and fibrils based on evidence from the ThT assay, AFM, DLS and GA derivative interaction modeling studies. Indeed, mixing GA with Aβ_1-42_ fibrils reduced Aβ_1-42_-induced calcium influx and protected cultured cortical neurons from toxicity.

The APP/PS1 transgenic mouse is an excellent model system to determine the development of cognitive impairments in mice over time. The 4-month-old APP/PS1 mice had very small amount of Aβ plaque deposition in the brain and mild cognitive deficits in the spatial reference and working memories which resemble those seen during the early stage of AD development in humans. In contrast, the 9-month-old APP/PS1 transgenic mice showed a large amount of Aβ_1-42_ plaques in the brain and severely compromised cognitive functions. This stage of disease development represents the late stage of AD seen in humans. GA treatment benefited both the early and late stage of APP/PS1 mice. The fact that GA significantly enhanced cognitive decline in APP/PS1 mice strongly suggests that GA is selective against Aβ_1-42_ and promises to serve as an effective drug candidate to delay and treat AD progression in humans.

Hippocampal LTP is considered one of the major cellular mechanisms underlying learning and memory. The 10-month-old APP/PS1 mice exhibited a significantly reduced LTP compared to those from the WT littermates and GA-treated mice (Fig 3). This difference demonstrated a strong correlation of GA in enhancing synaptic strength of the APP/PS1 mouse brain, and which also led us to further investigate the possible mechanisms. Two possible mechanisms were investigated which included: (1) GA disrupting Aβ_1-42_ aggregation *in vitro* and *in vivo;* and (2) GA inhibiting intracellular calcium influx to neurons. Multiple lines of evidence from the present study showed that GA inhibits Aβ_1-42_ forming aggregates both in a test tube and in the brain of APP/PS1 mouse brain. Oral administration of GA to the APP/PS1 AD mice effectively reduced the Aβ_1-42_ plaque size, rather than the Aβ_1-42_ plaque number, adding weight to the suggestion that GA alters Aβ_1-42_ aggregation to reduce its toxicity. This explains how GA treatment effectively reduced toxic Aβ_1-42_ load leading to long-term protection to synaptic functions.

Indeed, previous studies showed that GA was one of the most potent compounds in inhibiting kappa-casein fibril formation and stabilizing kappa-casein to prevent its aggregation^21.^ GA can also inhibit α-synuclein aggregation, which is another critical factor in the pathogenesis of AD ^38^. Based on the present atomic scale modeling studies and the ThT assay results, it is clear that the three hydroxyl groups of GA are essential in stabilizing the interaction of GA with Aβ_1-42_ aggregation. The -COOH group of GA appears to be essential in inhibiting the Aβ_1-42_ fibril formation. GA potentially disrupts the salt bridge between the residues of Lys28-Ala42 through non-covalent chemical bonding. Alternatively, oxidized GA and pyrogallol covalently interact with Aβ_1-42_ through Schiff-base formation between carbonyl group of the oxidized product and the ε-amino group of Lys16. Both interactions potentially alter the lateral association of layers of Aβ_1-42_ and prevent further Aβ_1-42_ aggregation.

NMDA receptor-mediated calcium influx is one of the major mechanisms of neuronal toxicity. Uncompetitive blockers of NMDA receptor-mediated intracellular calcium influx, such as memantine, confer potent protection against AD ^18,19^. Our data showed that GA could not completely inhibit glutamate and Aβ_1-42_ elicited intracellular calcium influx, indicating that GA’s protective effect was not through antagonizing these calcium receptors. Interestingly, pre-mixing GA with aggregated Aβ_1-42_ effectively reduced calcium influx confirming that GA works at the level of inhibiting Aβ aggregation.

There is ample circumstantial evidence to show that polyphenolic compounds from fruits, green tea and red wine can have beneficial effects in reducing the onset of AD. The present study represents a very detailed investigation into the impact and mechanism of such a compound. Future studies in usage of GA in a clinical setting are warranted.

## Acknowledgments

We thank SUSTech Animal Facility for the maintenance of transgenic mice. Financial supports to Dr. Sheng-Tao Hou came from the National Natural Science Foundation of China Grant (81571287), Shenzhen Science and Technology Innovation Committee Basic Science Research Grant (JCYJ20140417105742709, JCYJ20160301112230218); State Key

Laboratory of Neuroscience Open Competition Grant (SKLN-201403), SUSTech Peacock Program Start-up Fund (22/Y01226109) and SUSTech Brain Research Centre Fund. Financial support to Dr. Jing Xu and Dr. Chengqing Ning came from the National Natural Science Foundation of China (No. 21402082, No. 21772082 and No. 21702094), SZSTI (Pu20150267, Ji20170314 and Peacock Tech-Innovation 2018), and SZDRC (K16205905).

## Authors contributions

MY performed animal breeding, genotyping, behavioral studies, analyzed the data and wrote the first draft of the paper. XWC carried out neuronal cultures, AFM, ThT assay and calcium imaging studies; JHL did LTP recordings and brain slices; HC, LZ, and SY are responsible for the initial studies of Aβ aggregation, ThT assay, and dendritic studies; TMG was responsible for supervising the LTP studies, CQN and JX carried out molecular docking studies, and STH conceived the idea, provided financial support to the project, discussed the data and analyzed the results. STH revised the first draft and wrote the manuscript.

## SUPPLEMENTAL INFORMATION

**Fig. S1.**
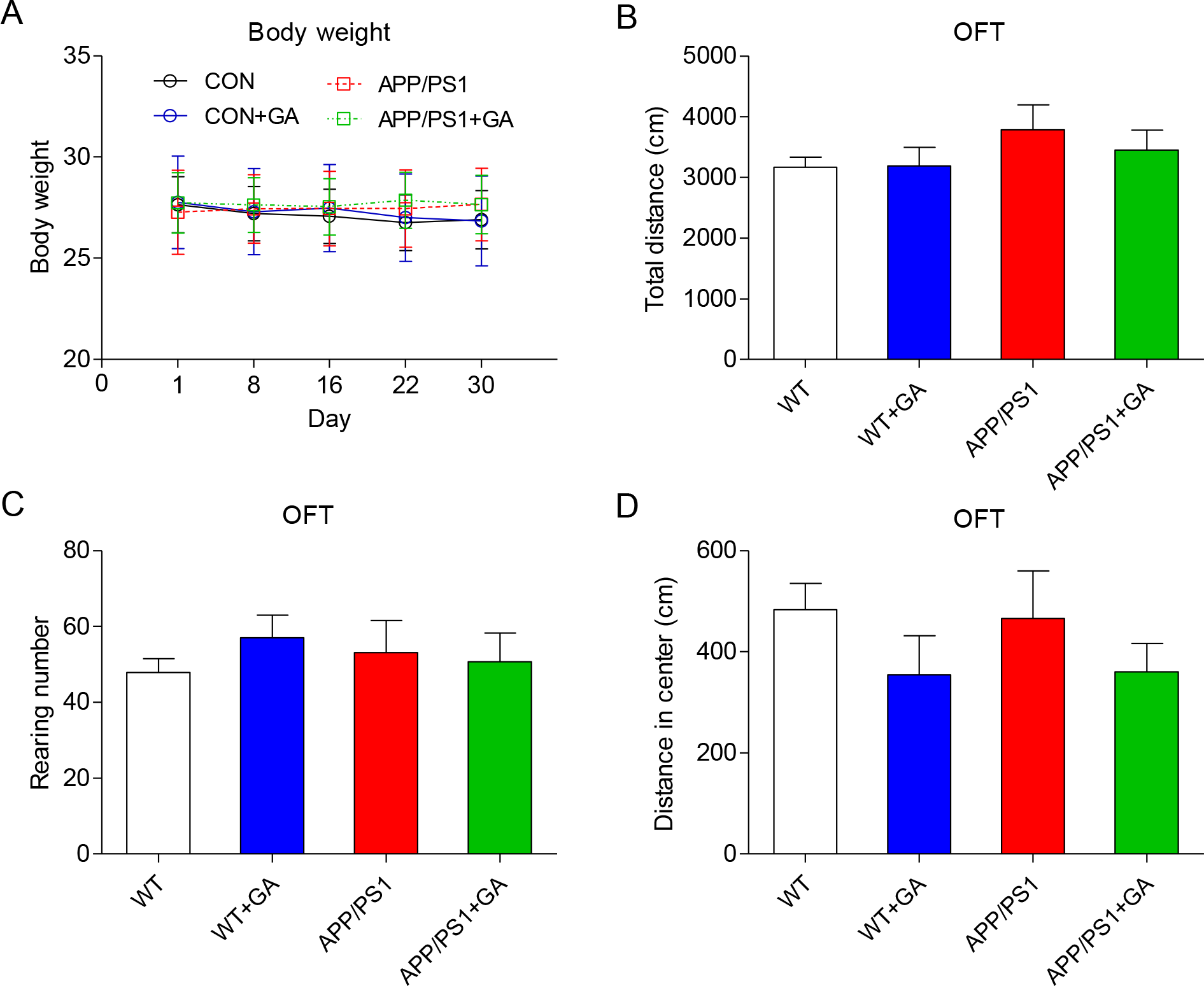
The body weight and spontaneous activity of five-month-old APP/PS1 transgenic mice were not affects by one month GA administration. (A) Body weight. (B) The whole traveled distance. (C) Rearing number. (D) The distance traveled in the center. Data represent mean ± SEM. Error bars indicate SEM. n = 6-10 mice in each group.

**Fig. S2.**
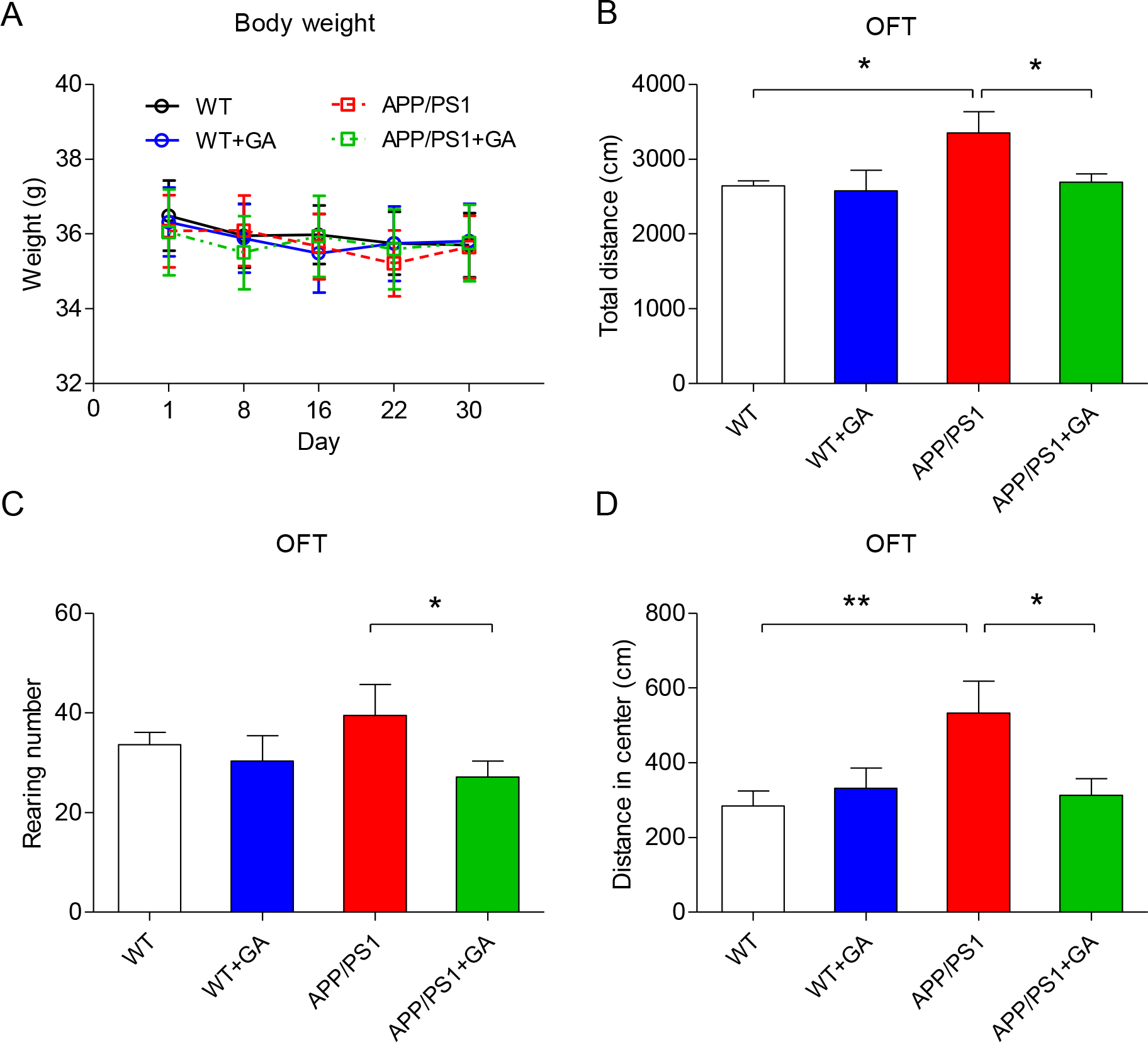
GA reduced the excessive spontaneous activity in ten-month-old APP/PS1 transgenic mice after one-month GA administration. (A) Body weight. (B) The whole traveled distance. (C) Rearing number. (D) The distance traveled in the center. Data represent mean ± SEM. Error bars indicate SEM. n = 6-10 mice in each group. * P < 0.05, * * P < 0.01.

**Fig. S3.**
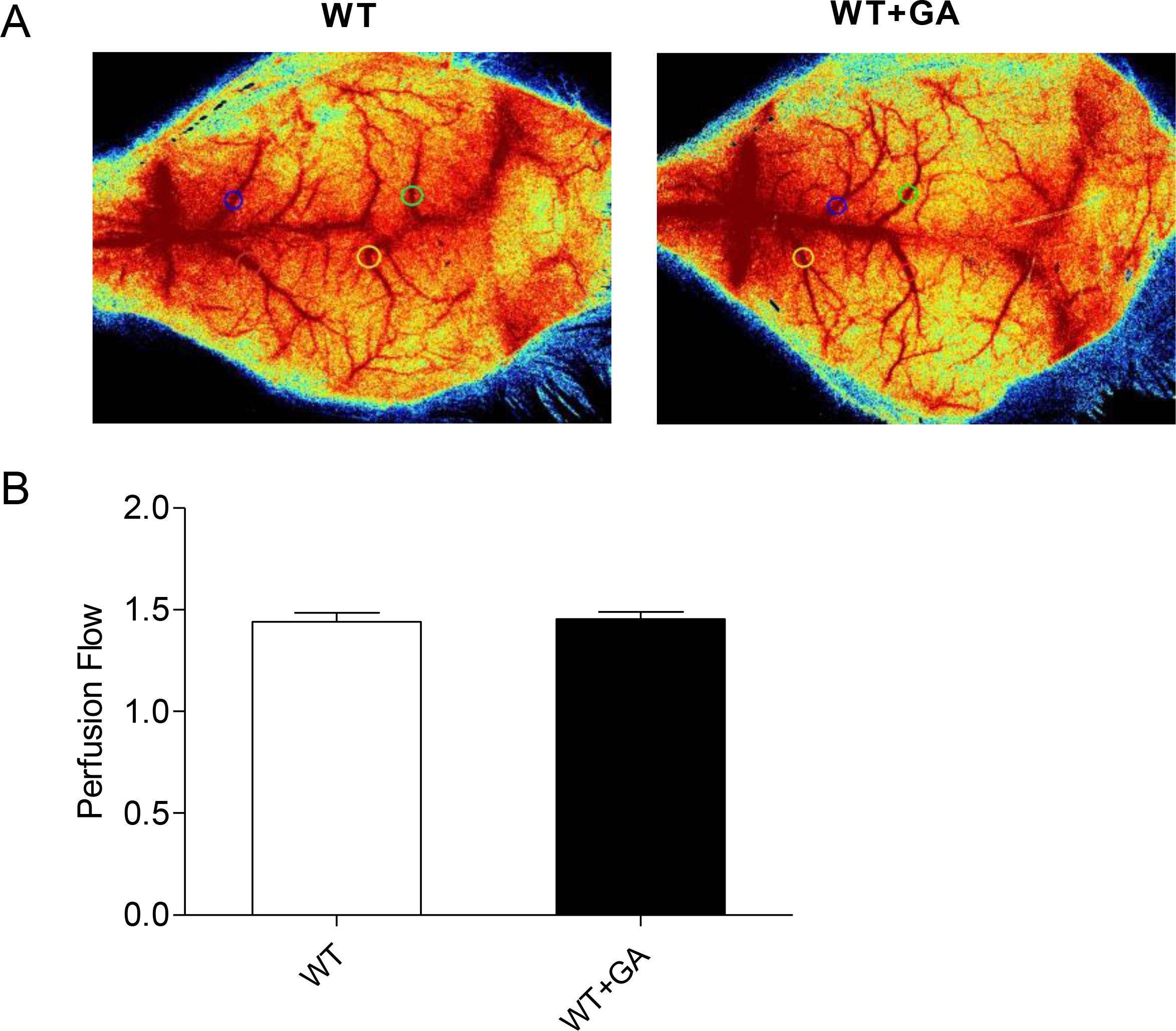
GA did not alter cerebral brain perfusion flow of mice, as determined by laser speckle contrast image (Moor FLPI-2: full-field laser perfusion imager). A: representative LSCI images and the setting of region of interesting (ROI, the circles of different colors). (B) The statistical result of perfusion flow in ROI, Data represent mean ± SEM. Error bars indicate SEM. n = 3 mice in each group.

